# TAM receptors mediate the Fpr2-driven pain resolution and fibrinolysis after nerve injury

**DOI:** 10.1101/2024.08.04.605987

**Authors:** Beate Hartmannsberger, Adel Ben-Kraiem, Sofia Kramer, Carolina Guidolin, Ida Kazerani, Kathrin Doppler, Dominique Thomas, Robert Gurke, Marco Sisignano, Pranav P. Kalelkar, Andrés J. García, Paula V. Monje, Asma Nusrat, Alexander Brack, Susanne M. Krug, Claudia Sommer, Heike L. Rittner

## Abstract

Nerve injury causes neuropathic pain and multilevel nerve barrier disruption. Nerve barriers consist of perineurial, endothelial, and myelin barriers. So far, it is unclear whether resealing nerve barriers fosters pain resolution and recovery. To this end, we analysed the nerve barrier property portfolio, pain behaviour battery, and lipidomics for precursors of specialized pro-resolving meditators (SPMs) and their receptors in chronic constriction injury of rat sciatic nerve to identify targets for pain resolution by resealing the selected nerve barriers. Of the three nerve barriers – perineurium, capillaries, and myelin – only capillary tightness specifically against larger molecules, such as fibrinogen, recuperated with pain resolution. Fibrinogen immunoreactivity was not only elevated in rats at the time of neuropathic pain but also in nerve biopsies from patients with (but not without) painful polyneuropathy indicating that sealing of the vascular barrier might be novel approach in pain treatment. 15R-HETE (hydroxyeicosatetraenoic acid), a precursor of aspirin-triggered lipoxin A4, were specifically upregulated at the beginning of pain resolution. Repeated local application of resolvin D1-laden nanoparticles or Fpr2 agonists sex-independently resulted in accelerated pain resolution and fibrinogen removal. Clearing macrophages (*Cd206)* and fibrinolytic pathways (*Plat)* were also induced while inflammation (*Tnfα)* and inflammasomes (*Nlrp3)* were unaffected by this treatment. Blocking TAM receptors (Tyro3, Axl, and Mer) and tyrosine kinase receptors linking haemostasis and inflammation completely inhibited all the effects. In summary, nanoparticles can be used as transporters for fleeting lipids, such as SPMs, and therefore expand the array of possible therapeutic agents. Thus, the Fpr2-Cd206-TAM receptor axis may be a suitable target for strengthening the capillary barrier, removing endoneurial fibrinogen, and boosting pain resolution in patients with chronic neuropathic pain.

## Introduction

Traumatic peripheral nerve injury elicits Wallerian degeneration and neuropathic pain succeeded by regeneration, with axonal regrowth, remyelination, and target organ innervation. While many patients recuperate, some fail to recover and suffer long lasting chronic pain. In the last few decades, research has provided valuable insights into chronification mechanisms, but healing and resolution of pain are less well-understood (Cohen et al., 2021). Regeneration failure includes incomplete restoration or miswiring of sensory and nociceptive fibres causing neuropathic pain (Gangadharan et al., 2022).

Immune cells are believed to play an important role in the switch to pain resolution which can occur independently of the recovery of tissue architecture and function (Sommer et al., 1993). While sensory or motor defects persist, the regeneration phase can emerge with mild or no pain. Immune cells control the pro-resolution switch, including macrophages and T cells, releasing anti-inflammatory cytokines (Fiore et al., 2023; Kavelaars & Heijnen, 2021; Sim et al., 2023), specific subtypes of spinal microglia (Donovan et al., 2024) and mitochondria or extracellular vesicles from anti-inflammatory macrophages (van der Vlist et al., 2022). These highly-regulated processes balance the switch for chronification or resolution (Parisien et al., 2022). However, the temporal course in nerve tissues is not thoroughly understood.

Nerve barriers are composed of perineurial, endothelial, and Schwann cells that shield the delicate nerve milieu from external stimuli (Moreau et al., 2016; Napoli et al., 2012; Reinhold et al., 2023). Under homeostatic conditions, nerve barriers ensure proper nerve function and conduction through tight junction proteins, such as claudins, pericytes, and endoneurial macrophages (Malong et al., 2023; Reinhold et al., 2023). Endoneurial oedema represents one of the earliest signs of nerve damage and barrier leakage in patients with inflammatory and non-inflammatory polyneuropathy and preclinical models (Sommer et al., 1993; Üçeyler et al., 2016), which manifests as endoneurial deposition of plasma proteins such as fibrinogen (Schenone et al., 1988), and loss of myelin barrier proteins (Chen et al., 2023).

After nerve injury, barrier breakdown – evident as tight junction protein loss and immune cell infiltration – facilitates the proper removal of myelin debris. Following successful clearance, nerve barriers need to be resealed to protect the regenerated sensitive structures for the proper nerve. The proposed factors that promote barrier resealing are TAM receptors (Tyro3, Axl, and Mer), which are part of a group of receptor tyrosine kinases that control efferocytosis in both macrophages and Schwann cells (Brosius Lutz et al., 2017; Elliott et al., 2017). TAM receptors mediate the resolution of inflammation and promote the production and synthesis of anti-inflammatory and specialized pro-resolving mediators (Vago et al., 2021). These SPMs – other factors for barrier resealing – derived from omega-3 and omega-6 polyunsaturated fatty acids, actively resolve inflammation (Tao et al., 2020). SPMs such as maresins, protectins, and resolvins activate seven different cell surface G-protein coupled receptors (GPCRs), including formyl peptide receptor 2 (FPR2) and G protein-coupled receptor 37-like 1 (GPR37L1), thereby ameliorating inflammatory and neuropathic pain (Bang et al., 2024; Su et al., 2023), endothelial cell activation, and vascular smooth muscle cell remodelling (Díaz Del Campo et al., 2022).

In this study, we explored the interplay between the nervous system, barrier function, and the immune system to create a temporal signature of pain resolution after traumatic nerve injury. Specifically, we employed in-depth lipidomics, barrier mapping, and pathway analysis and therapeutically used SPM-laden nanoparticles to foster pain resolution.

## Results

### 1. Resolution of thermal and mechanical hyperalgesia in chronic constriction injury from 4-6 weeks

We characterized the pain course using reflexive and non-reflexive tests over a period of nine weeks to determine the period of resolution of neuropathic pain after chronic constriction injury (CCI) in Wistar rats **(Fig. 1A)**. Mechanical and thermal thresholds decreased until week 4 after CCI compared with sham animals and completely recovered at week 6 in both males (**Fig. 1B, C**) and females (**SFig. 1**). Assessment of the mean print area and the stand time in gait analysis can serve as a measure of pain (Heinzel et al., 2020). The gait pattern drastically changed after CCI, while the gait of the sham-operated rats remained unaffected (**Fig. 1D, E**). Gait recovery took a similar but delayed course compared with reflexive tests. The stand time did not completely return to baseline levels after nine weeks, which is thought to result from nerve damage rather than pain. Despite their impaired gait, CCI animals demonstrated normal physical activity throughout the entire time course, using a voluntary running wheel (**Fig. 1F, G**).

**Figure 1.**
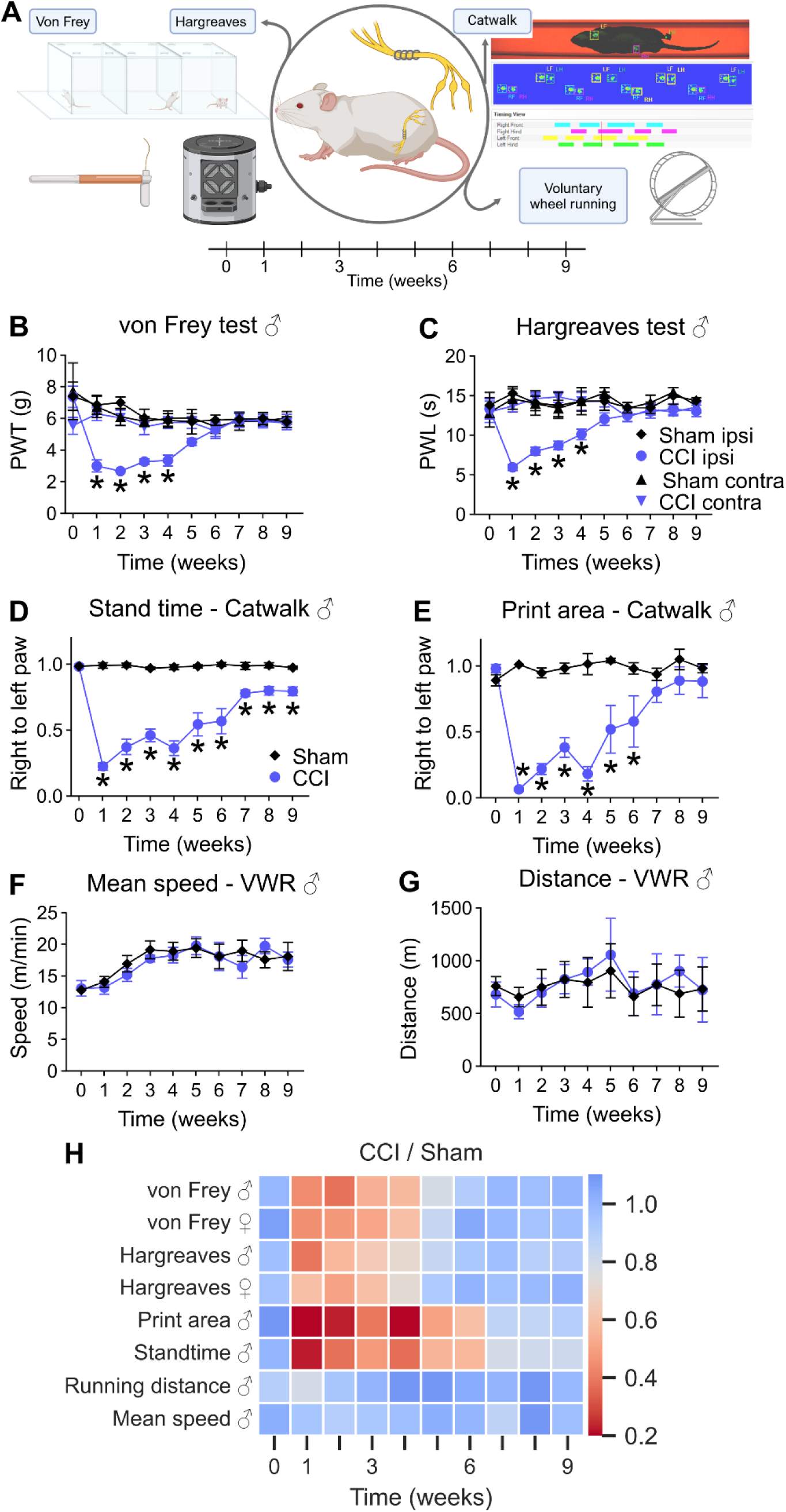
Neuropathic pain resolves 6 weeks after nerve injury. **(A)** Schematic illustration of the experimental setup: Male and female Wistar rats underwent CCI or sham surgery and were examined using the displayed tests throughout 9 weeks. **(B)** Mechanical allodynia and **(C)** thermal hypersensitivity were assessed weekly (n = 6-8). **(D)** The print area and **(E)** standtime presented as ratios of right to left hindpaws were recorded using the Catwalk gait analysis system (n = 6-7). **(F)** Covered distance and **(G)** mean speed of CCI and sham animals were recorded with voluntary running wheels (n = 8). **(H)** Summary of all behavioural tests including von Frey and Hargreaves from female rats. Heatmap shows the ratios between arithmetic means of CCI and sham groups. Data in graphs are shown as mean ± SEM, * p<0.05. Two-way repeated measures ANOVA with Tukey’s multiple comparison. PWT: paw withdrawal threshold; PWL: paw withdrawal latency; CCI: chronic constriction injury; VWR: voluntary wheel running.

Based on the curves for mechanical and thermal hyperalgesia and footprint analysis, pain resolution started from week 3 to week 4 after CCI, whereas the end of the resolution was evident at week 6 (**Fig. 1H**). Thus, we defined week 1 after surgery as maximal hypersensitivity, week 3 as the flexion point of resolution, week 6 as the end of pain resolution, and week 9 as the nerve regeneration phase.

### 2. Capillaries reseal for fibrinogen in parallel to pain resolution

To determine whether pain resolution coincided with the resealing of any of the nerve barriers, we performed a battery of permeability tests and tight junction protein analysis **(Fig. 2A)**. Hyperpermeability of the perineurium persisted for fluorescein (376 Da) and Evan’s Blue albumin (EBA, 69 kDa) for up to nine weeks after CCI (**Fig. 2B-D**). To assess the state of the myelin barrier in the proximal, non-degenerating part of the nerve, we immersed desheathed nerve fibers in fluorescein isothiocyanate (FITC)-dextran (70 kDa). Dye leakage through the myelin barrier was similar after nerve injury compared with that in the sham group (**Fig. 2E, F**). Perineurium breakdown was reflected by the low levels of *Cldn1* and *Tjp1* **(Fig. 2G, H)** in the ligated region, whereas *Cldn19* expression was heavily reduced **(Fig. 2I)** according to *Mpz* levels **(SFig. 2A)**.

**Figure 2.**
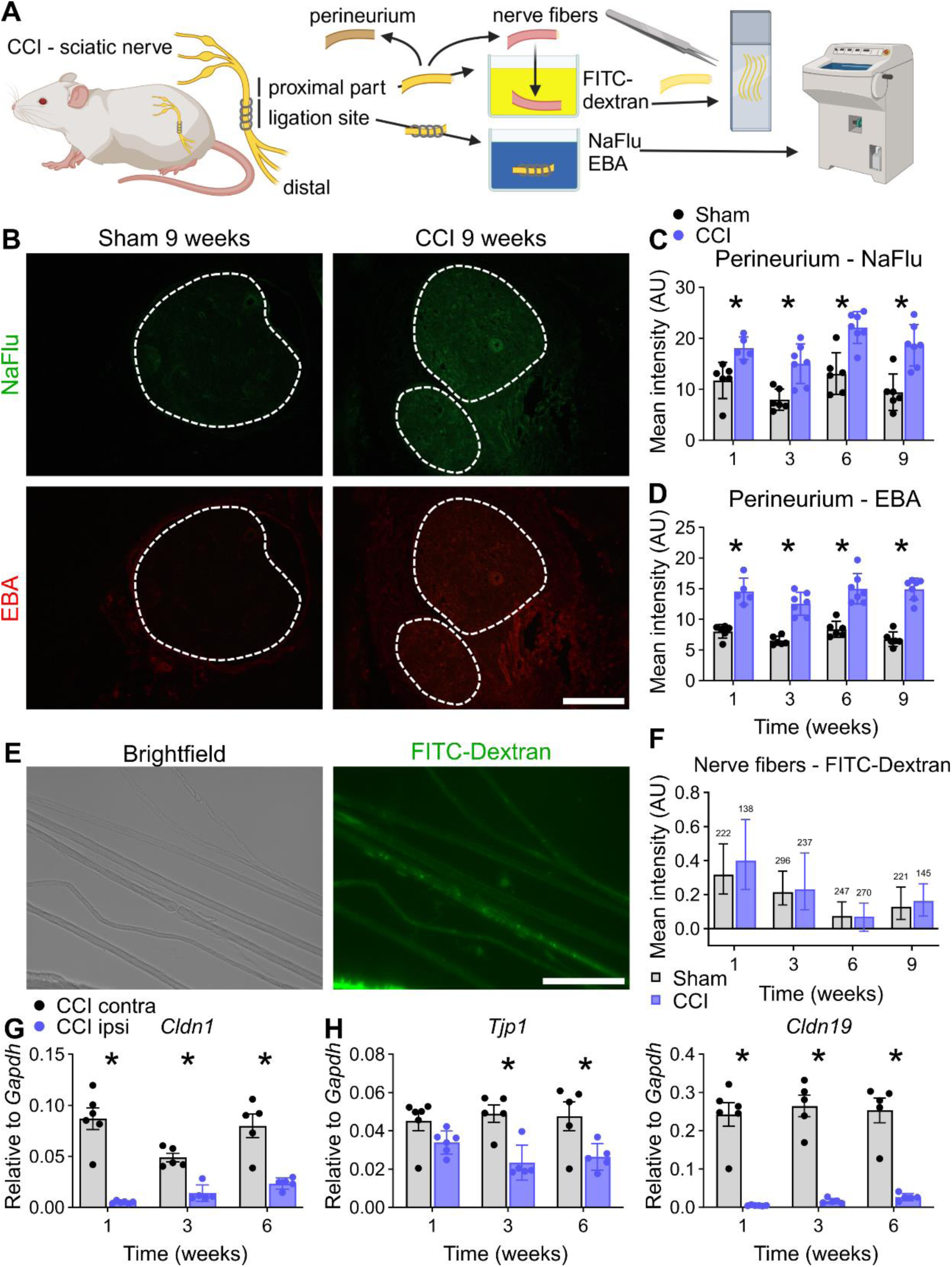
The perineurial barrier remains open up to 9 weeks while myelin barrier stays intact. **(A)** Depiction of two *ex vivo* immersion techniques for analyzing the perineurial and myelin barriers CCI or sham. **(B)** Representative images of sciatic nerve cross sections and **(C, D)** quantification of endoneurial fluorescence intensity of nerves immersed in NaFlu (376 Da) and Evans blue albumin (EBA, 68 kDA) *ex vivo*. Dashed lines indicate the endoneurial regions. Scale bar: 300 µm. **(E)** Representative brightfield and fluorescence images of teased nerve fibers from the proximal part of the sciatic nerve after immersion with 70 kDa FITC-dextran. Scale bar: 100 µm. **(F)** Quantification of the fluorescence intensity measured in the internodal regions of teased fibers. (n = 138-296 from 5-6 animals per group, pairwise Mann-Whitney-U-tests). Relative gene expression of **(G)** *Cldn1*, **(H)** *Tjp1*, and **(I)** *Cldn19* in the ligation parts of ipsilateral and contralateral sciatic nerves after 1, 3, and 6 weeks after CCI (n = 5-6). All data are shown as mean ± SEM, * p<0.05 compared to control at the indicated time points, two-way ANOVA with Šidák’s multiple comparisons unless indicated otherwise.

The integrity of the intraneural blood vessel barrier was compromised from weeks 1-6 for EBA and reestablished at week 9 (**Fig. 3A-D**). Blood vessels leaked at the ligation and distal parts for the intravenously-injected EBA, whereas the blood-nerve barrier (BNB) in the proximal parts further away from the ligations was unaffected. Accordingly, *Cldn5* was reduced in the ligated region (**Fig. 3H**). *Cldn12*, another tight junction protein, was also reduced and did not recover with pain resolution **(SFig. 2B)**.

**Figure 3.**
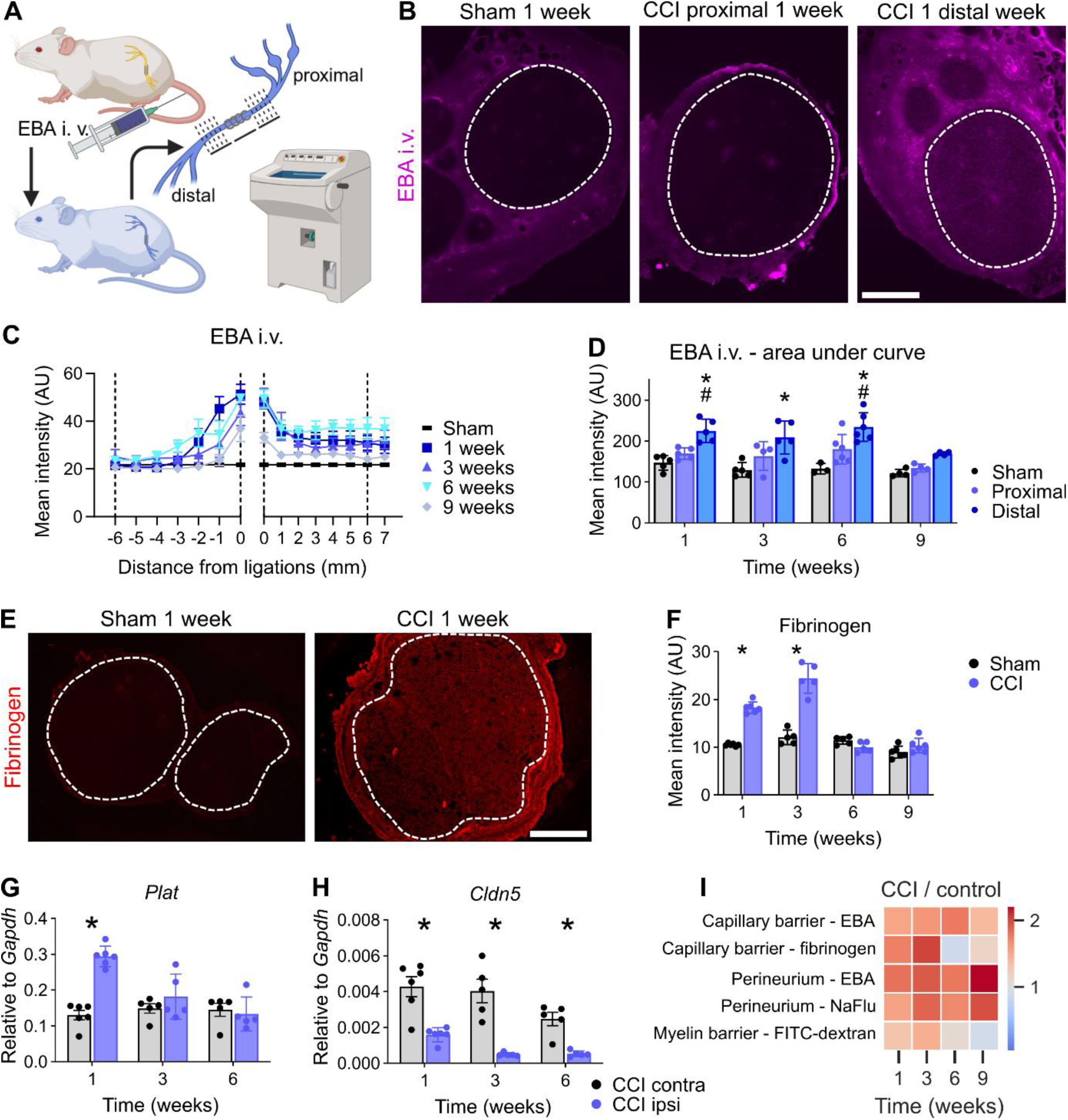
Capillary barrier reseals specifically for fibrinogen in parallel with neuropathic pain. **(A)** Schematic illustration of the capillary permeability test: Evans blue albumin (EBA, 68 kDA) was intravenously injected. Cross sections at distinct positions distal and proximal from the ligations were collected and analysed. **(B)** Representative images of sciatic nerve cross sections. The dashed lines indicate the endoneurial regions. Scale bar: 300 µm. **(C)** Depiction of the spatial and temporal EBA extravasation into the endoneurium through endoneurial vessels. Sham values were averaged and plotted on each position as reference. Dashed lines mark the positions that were used for the calculations for the areas under the curve in **(D)**: Quantification analysis of the areas under the curves distal and proximal from 0 mm to 6 mm from the ligations were plotted (n = 3-6). *: distal vs. sham; #: distal vs. proximal p<0.05 two-way ANOVA with Bonferroni’s multiple comparisons test. **(E)** Representative images of fibrinogen immunoreactivtiy in the sciatic nerve 1 week after surgery. The dashed lines indicate the endoneurial region. **(F)** Quantification of the intensity of fibrinogen immunoreactivity within the endoneurial regions (n = 5-6). Scale bar: 300 µm. Relative gene expression of **(G)** *Plat* and **(H)** *Cldn5* in sciatic nerves after CCI (n = 5-6). **(I)** Summary of all permeability tests. The heatmap displays the ratios between arithmetic means of CCI and sham groups. All data are shown as mean ± SEM, * p<0.05 two-way ANOVA with Šidák’s multiple comparisons.

Endoneurial leakage of blood proteins is a well-known hallmark of polyneuropathy (Schenone et al., 1988). Fibrinogen (340 kDa) is a blood coagulation factor, produced in the liver, and polymerizes to fibrin upon tissue damage. Tissue plasminogen activator (tPA; gene: *Plat*) activates plasminogen to degrade fibrin. In injured nerves, fibrinogen accumulated endoneurially and *Plat* was upregulated until 3 weeks (**Fig. 3E-G**). When hypersensitivity was resolved, both fibrinogen and *Plat* returned to sham levels, indicating complete recovery of the BNB in terms of fibrinogen leakage. The capillary barrier was impermeable to fibrinogen at the time of complete pain resolution.

Endoneurial fibrinogen correlated with the pain course in the analysis of the barrier portfolio (**Fig. 3I**), indicating that the presence of fibrinogen, a pro-inflammatory and pro-nociceptive molecule (Akassoglou et al., 2000; Lim et al., 2014) that penetrates the capillary BNB into the endoneurium, might explain the pathophysiology of the switch to pain resolution.

### 3. Human nerves contained more endoneurial fibrinogen in painful neuropathy

To validate our preclinical findings in a translational approach, we analysed sural nerve biopsies of patients with different polyneuropathies (**Tab. 1**). Patients were 69 ± 7.8 years, mainly male (10 males, 6 females) with a disease duration of 52 ± 44.1 months and a clinical severity of the neuropathy of 2 ± 1.2 (overall disability sum score (ODSS) score, 0 = no sign of disability to 12 = most severe disability). The demyelinating and axonal neuropathies were almost evenly distributed. Half of the patients (8 of 16) had pain at the time of nerve biopsy while the other half did not. Follow-up information was available from only six patients. Three of them had pain, of which only one was painless several years later. We immunostained the nerves and compared the fibrinogen and tPA signal intensities (**Fig. 4A, C**). Patients with painful polyneuropathy showed higher fibrinogen immunoreactivity in the endoneurium, whereas the tPA levels did not differ (**Fig. 4B, D**). These results suggest that fibrinogen may also play a role in the persistence of pain in human patients with painful neuropathy.

**Figure 4:**
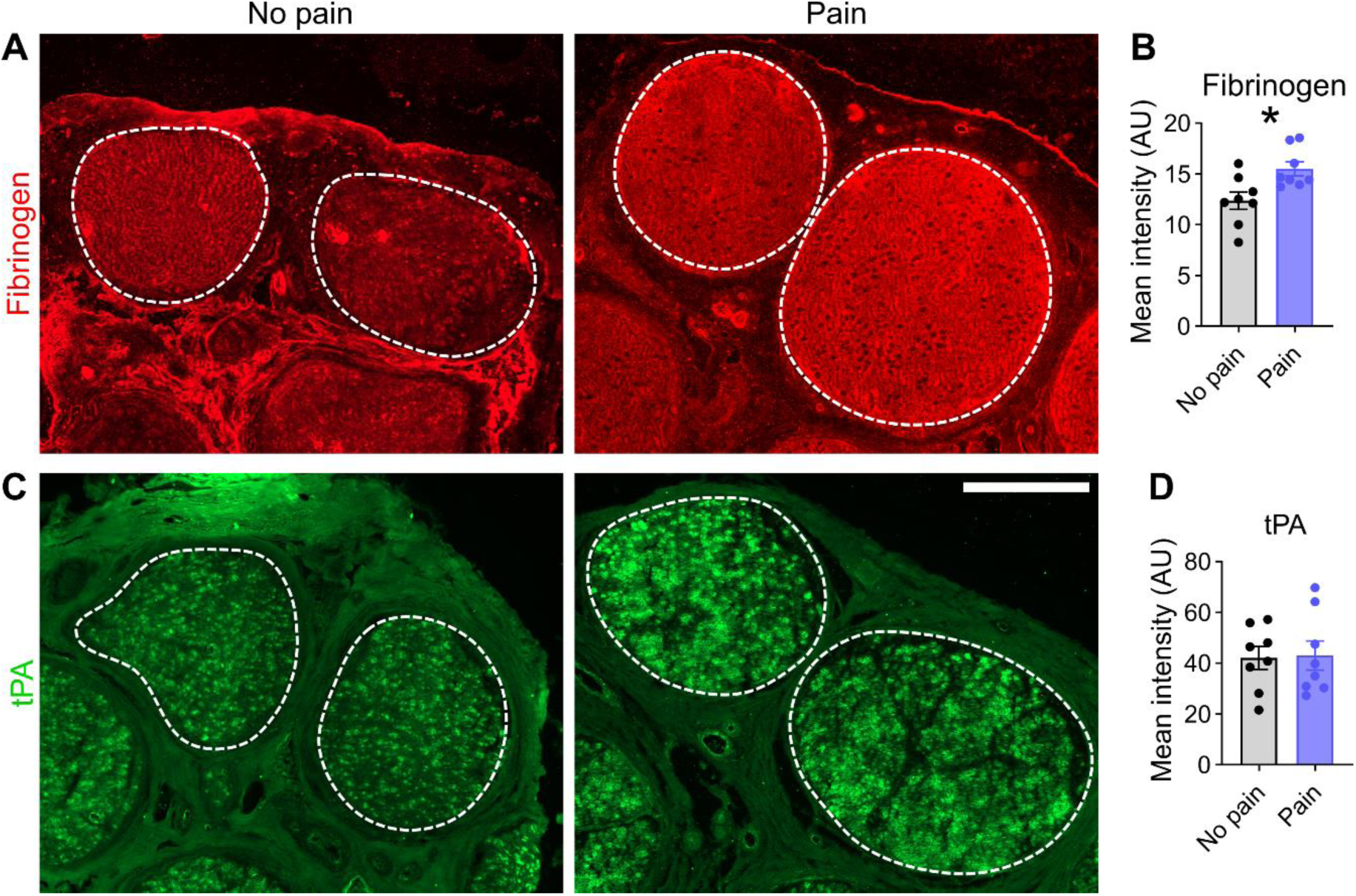
More endoneurial fibrinogen in sural nerve biopsies from patients with painful polyneuropathies. Sural nerve biopsies from patients were immunofluorescently stained. Representative images of human sural nerves stained for **(A)** fibrinogen and **(C)** tissue plasminogen activator (tPA). The dashed lines indicate the endoneurial regions. Scale bar: 300 µm. **(B, D)** Quantifiactions of signal intensity of fibrinogen and tPA. Patients were grouped by pain occurence (n=16, Student’s t-tests with Welch’s correction).

**Table 1.** Patient demographic and clinical characteristics of painful and non painful neuropathies.

| Age | Sex | Diagnosis | Duration (months) | ODSS | Pain | Histology |
| --- | --- | --- | --- | --- | --- | --- |
| 77 | f | CIAP | 48 | 2 | yes | mixed |
| 72 | m | CIDP/atypical | 12 | 2 | yes | demyelinating |
| 71 | m | CIAP | 48 | 2 | yes | axonal |
| 67 | f | idiopathic PNP | 24 | 3 | yes | axonal |
| 73 | m | CIDP/atypical | 36 | 6 | yes | axonal |
| 57 | f | idiopathic PNP | 6 | 1 | yes | axonal |
| 73 | m | CIDP/atypical | 9 | 3 | yes | demyelinating |
| 62 | f | idiopathic PNP | 66 | 1 | yes | axonal |
| 64 | m | CMT | 36 | 2 | no | axonal |
| 59 | f | CIDP | 120 | 3 | no | demyelinating |
| 71 | m | idiopathic PNP | 120 | 2 | no | demyelinating |
| 66 | f | idiopathic PNP | 120 | 2 | no | demyelinating |
| 76 | m | CIDP | 12 | 2 | no | demyelinating |
| 53 | m | CMT | 48 | 3 | no | demyelinating |
| 80 | m | CIDP/atypical | 7 | 2 | no | axonal |
| 76 | m | CIAP | 120 | 1 | no | axonal |

### 4. Lipidomics: 15R-HETE culminates selectively at the switch for pain resolution

To identify the lipids driving the lipid mediator class switch from pro-inflammatory to pro-resolving and their respective receptors, we performed lipidomic analysis by LC-MSMS and qPCR of the sciatic nerve **(Fig. 5A)**. We analysed 34 lipids derived from arachidonic acid (AA), docosahexaenoic acid (DHA), and eicosapentaenoic acid (EPA), and detected 11 of them. Although pro-inflammatory prostaglandin E2 (PGE2) and thromboxane B2 (TXB2) were more abundant after injury, prostaglandin D2 (PGD2) levels were reduced. Among the three hydroxyeicosatetraenoic acid (HETE) precursor lipoxins (5S-, 15S-, and 15R-HETE), only 15R-HETE selectively increased at the beginning of pain resolution (**Fig. 5B**). Specialized pro-resolving mediators, such as neuroprotection D1 (NPD1), resolvin D1 (RvD1), resolving D2 (RvD2), and maresin 1 (MaR1), could not be detected. Other lipids from pro-resolving synthesis pathways, such as 17S-HDHA (precursor of RvD1 and 2) and 14S-HDHA (derived from DHA), showed higher concentrations, but not selectively, at 3 weeks (**Fig. 5B**). Lipids derived from EPA (18R-HEPE) were not detected.

**Figure 5.**
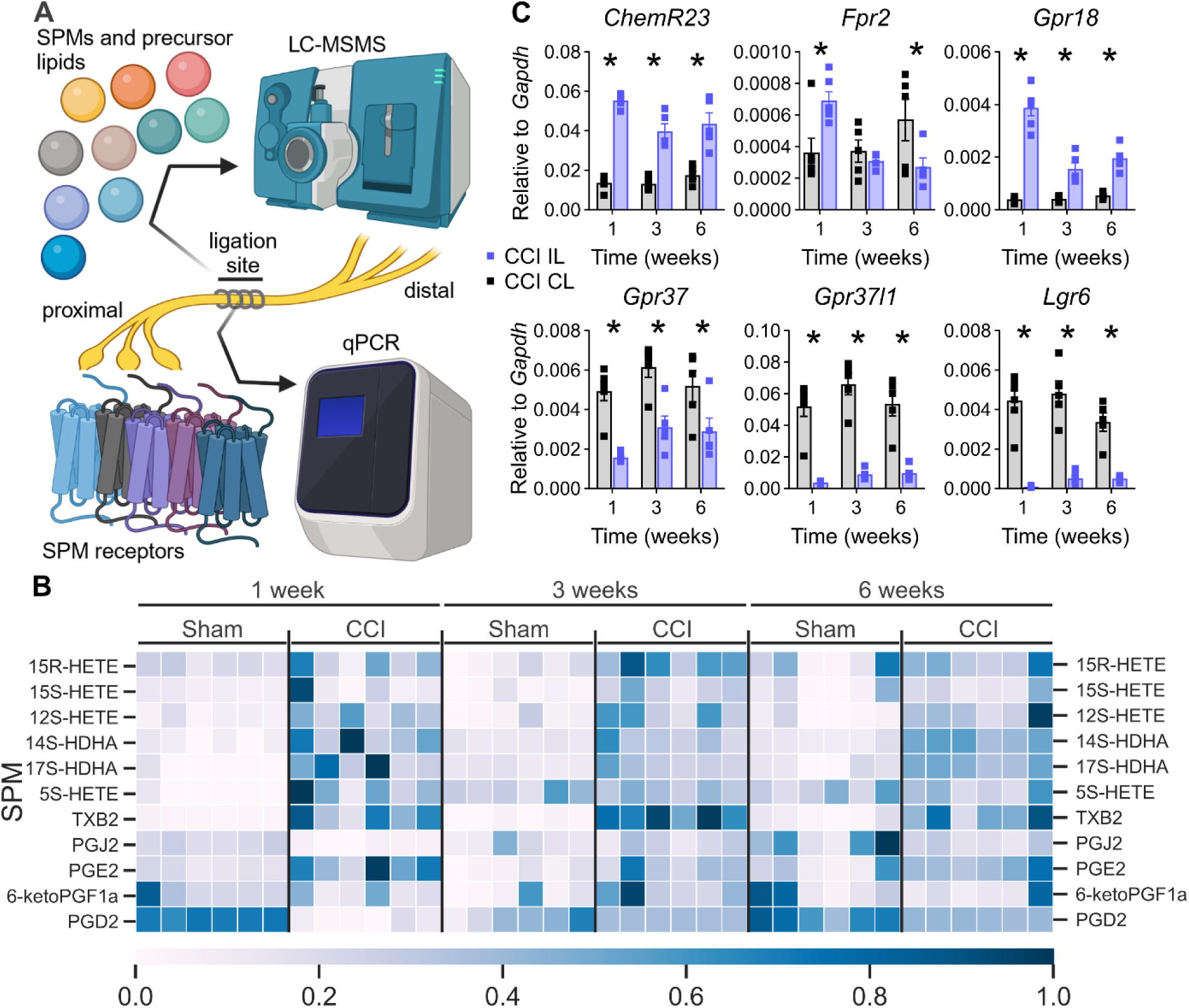
Unbiased lipidomic analysis selectively identified higher levels of the LXA4 precursor 15R-HETE at 3 weeks initiating the resolution phase. **(A)** Arachidonic, docosahexaenoic, and eicosapentaenoic acids-derived metabolites were measured by LC-MSMS in sciatic nerve, and their cognate receptors were assessed using qPCR. **(B)** Relative levels of the 11 detected lipid metabolites are depicted. Only 15R-HETE had a significant upregulation in CCI specifically at 3 weeks compared with sham. **(C)** Relative mRNA expression of the respective SPM receptor after CCI; CL: contralateral; IL: ipsilateral. All data are shown as mean ± SEM, n = 5-6, two-way ANOVA with Šidák’s multiple comparisons. * p<0.05.

To further evaluate the potential of the aspirin-triggered lipoxin A4 (AT-LXA4) precursor 15R-HETE to resolve pain, we determined the mRNA expression of its receptor, formyl peptide receptor 2 (*Fpr2*), and other known SPM receptors *Gpr18, Chem23r, Gpr37*, and *Lgr6* (Fig. 4C)*. Gpr18* and *ChemR23* were upregulated, whereas *Lgr6*, *Gpr37, and Gpr37l1* were downregulated. *Fpr2* was not abundant but was increased at 1 week and decreased 6 weeks after injury (**Fig. 5C**) but is known to be expressed in rat macrophages (Rittner et al., 2009). To determine the abundance of these receptors in Schwann cells, the most abundant cell type in nerve fibres, we measured them in a primary Schwann cell culture. Except for *Gpr37* (C_T_ value: 28.9) and *ChemR23* (C_T_ value: 31.7), all C_T_ values were above 32 and were not relevantly expressed. Since Gpr37 is specifically expressed in promyelinating Schwann cells during myelination, we hypothesized that *Gpr37* is the only suitable candidate for Schwann cell differentiation. cAMP stimulation induced rapid upregulation of *Gpr37* after 24 h but did not increase *Gpr37l1* and *Mpz* expression after 5 days (**SFig. 3**). This indicates that, in our experiment, *Gpr37* might be a suitable target after pain resolution has occurred.

### 5. Fpr2 activation by BML-111 accelerates pain resolution and degradation of fibrinogen

Since we identified the AT-LXA4 precursor at the pain resolution switch, we further investigated Fpr2 at 2.5 weeks preceding “natural” pain resolution. To specifically activate Fpr2 in the sciatic nerve, we applied the Fpr2 agonist BML-111 perineurially twice daily for 1 week and tested its analgesic effects using the von Frey and Hargreaves tests (**Fig. 6A**). The analgesic effect of 500 nmol BML-111 arose from 1-3 h after injection and vanished after 6h (**Fig. 6B**). Therefore, we decided to administer a second dose after the behavioural test. Cumulative analgesic effects (measured before the first injections of the day) appeared on the third day of injections and gradually increased during the 7-days injection period. While mechanical hypersensitivity significantly improved on day 6, thermal hypersensitivity significantly improved on day 2 and later reached the contralateral paw withdrawal thresholds (**Fig. 6C, D**). After 1 week of treatment, fibrinogen deposition was reduced in BML-111-treated nerves compared to that in vehicle-treated nerves (**Fig. 6E, F**). Elevated *Plat* levels in BML-111-treated nerves suggest that fibrinolysis was still in process (**Fig. 6G**) and was not terminated, as observed in CCI animals after 6 weeks (**Fig. 3G**).

**Figure 6.**
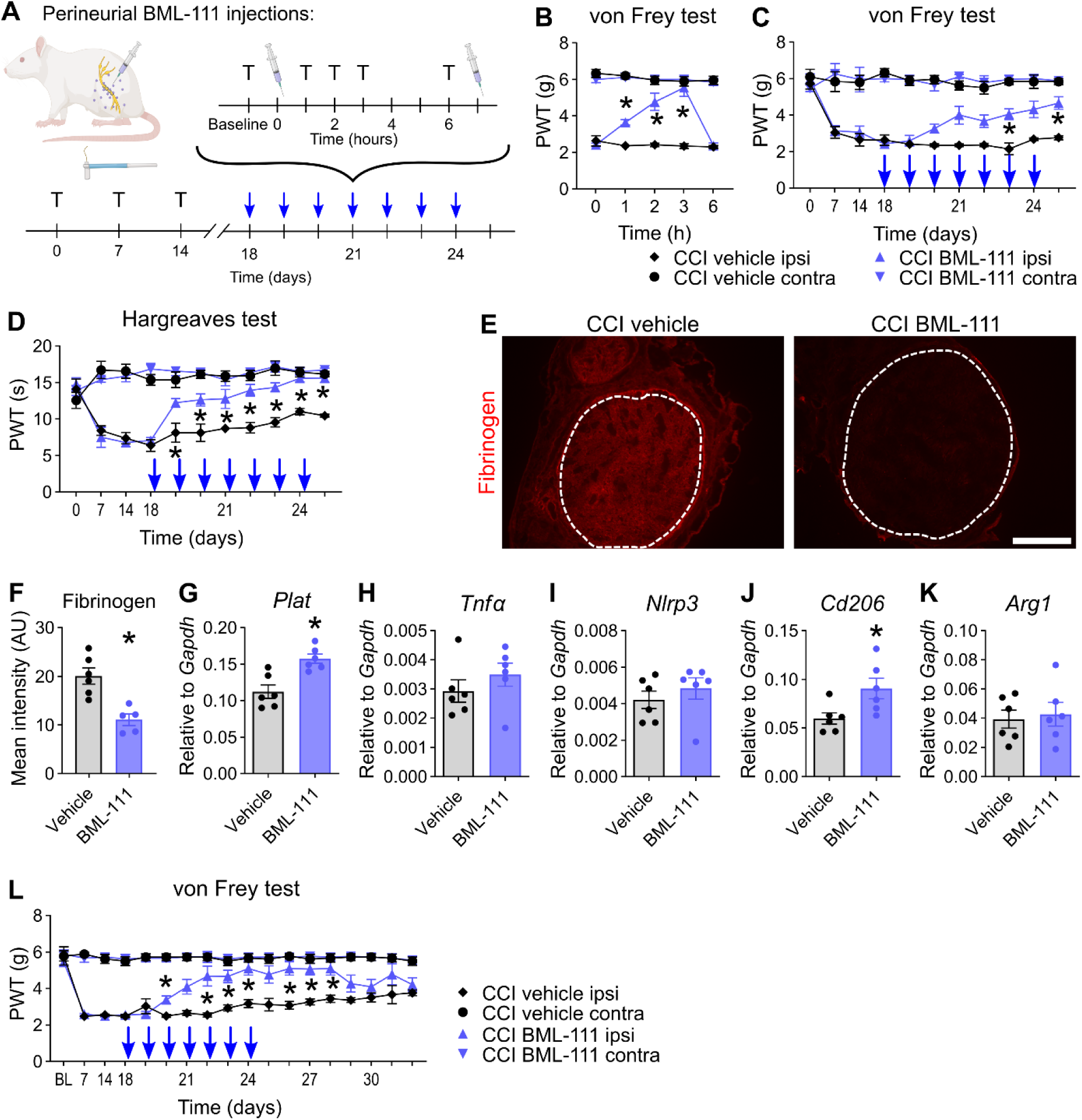
Local application of the Fpr2 agonist BML-111 initiates pain resolution, macrophage polarization, and fibrinogen clearance. **(A)** CCI-operated animals received a perineurial injection of BML-111 or vehicle (PBS). Animals were retested with Hargreaves and von Frey at 1, 2, 3, and 6 h. As soon as reflexive tests at 6 h were finished, the animals received a second injection from day 18 to 24, and were sacrificed at day 25. Blue arrows indicate the daily injection/test program. T: reflexive testing; syringe: injection. **(B)** Mechanical allodynia was measured after the first BML-111 injection on day 18 (n = 6, two-way repeated measures ANOVA with Tukey’s multiple comparisons). Paw withdrawal thresholds for **(C)** mechanical and **(D)** thermal hypersensitivities of baseline measurements show an accumulative effect of perineurial BML-111 (n= 5-6, two-way repeated measures ANOVA with Tukey’s multiple comparisons). **(E, F)** Representative images and quantification of fibrinogen immunostaining in the sciatic nerve after BML-111 injections. The dashed lines indicate the endoneurial region. Scale bar: 300 µm. (n = 6, Student’s t-tests with Welch’s correction). **(G-K)** Relative mRNA expression of *Tnfα*, *Arg1*, *Cd206*, *Nlrp3*, and *Plat* after BML-111 injections (n = 6, Student’s t-tests with Welch’s correction). **(L)** Mechanical allodynia was assessed daily for one week after injections were discontinued. (n = 6, two-way repeated measures ANOVA with Tukey’s multiple comparisons). All data are shown as mean ± SEM, *: p<0.05.

Fpr2 inhibits pro-inflammatory and pronociceptive genes such as *Tnfα*, *Nlrp3* and promotes anti-inflammatory macrophages expressing *Arg1* and *Cd206* (Cao et al., 2018; Yuan et al., 2022). In our system, only *Cd206* expression was increased (**Fig. 6H-K**) highlighting the role of endoneurial fibrinogen. Over the course of 6 weeks, the increased abundance of *Tnfα*, which is known to induce pain, in the CCI nerve compared to sham suggests that *Tnfα* might not be critically involved in pain resolution (**SFig. 4**). We checked whether *Cd206* was upregulated in the phase of pain resolution; however, *Cd206* levels mirrored the *Plat* levels, while *Cd68* remained stably elevated after 6 weeks (**SFig. 4**).

To determine whether the activation of Fpr2 induces permanent or irreversible pain relief, we tested the mechanical allodynia of BML-111-administered animals for one week seven days after injections. After the last BML-111 treatment, mechanical thresholds remained stable for three days and re-approached the levels of vehicle animals, which showed slowly resolving pain (**Fig. 6L**).

### 6. RvD1 nanoparticles also accelerate resolution in male and female rats

Analgesia by BML-111 ceased rapidly, and SPMs were even shorter lasting. To evaluate SPMs as therapeutic long-lasting Fpr2 agonists, we applied RvD1-laden polymeric nanoparticles (RvD1-NPs, **Fig. 7A**). Nanoparticles released RvD1 with maximum concentrations between 3 and 24 h (**Fig. 7B**). Based on these kinetics, RvD1-NPs only needed to be applied once daily to ensure continuous RvD1 supply. RvD1-NPs elicited analgesic effects on mechanical allodynia (**Fig. 7C, D**). RvD1-NPs caused fibrinogen degradation in the endoneurium and a trend for increased *Plat* (**Fig. 7E-G**). Similar to BML-111-treated animals, *Tnfα*, *Nlrp3,* and *Arg1* levels did not change after RvD1-NP injection, and *Cd206* was upregulated (**Fig. 7H-K**). Similar effects were observed after RvD1-NP treatment in female rats, suggesting a similar mechanism in male and female rats (**Suppl. Fig. 5**).

**Figure 7.**
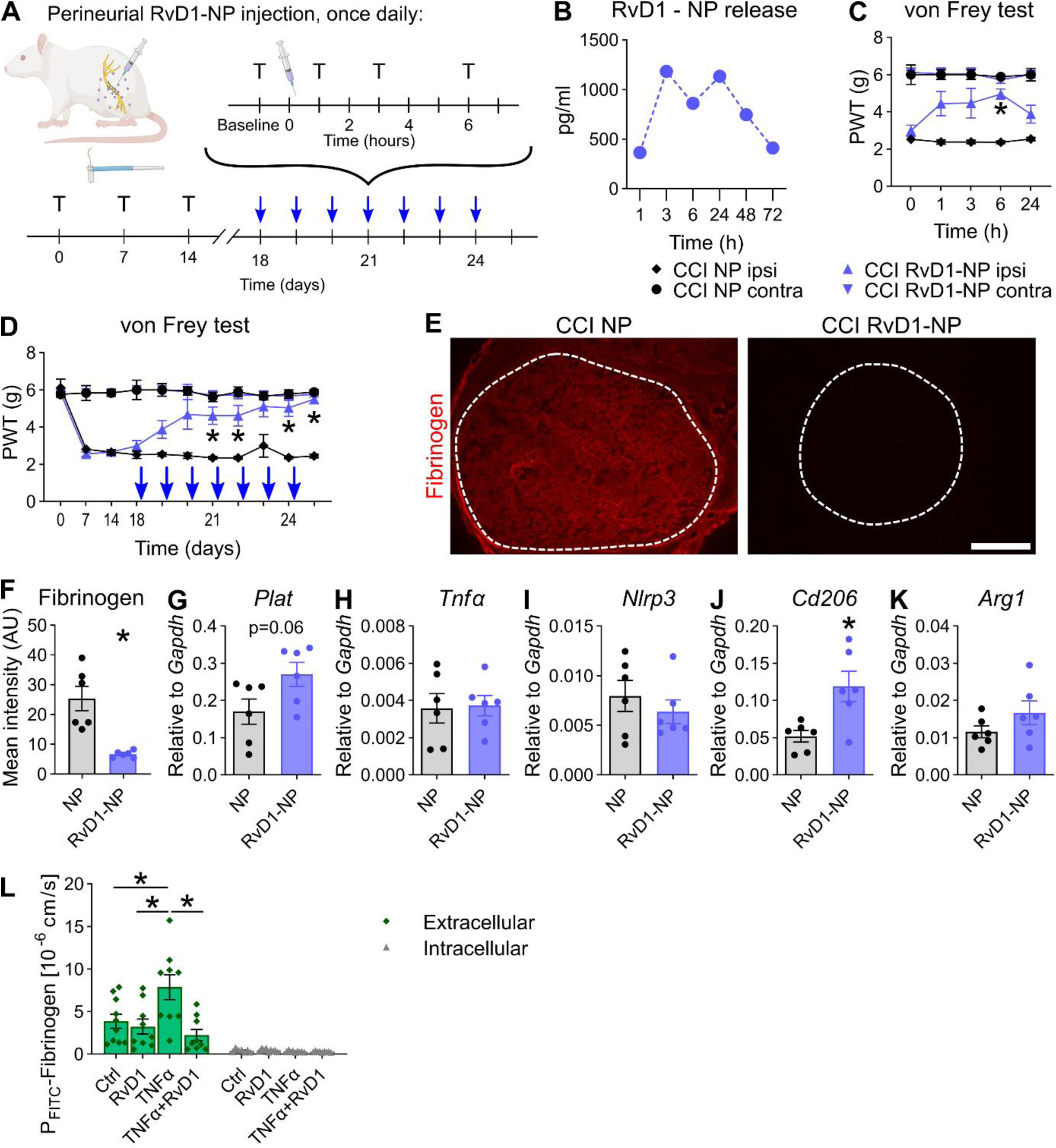
Locally applied RvD1-laden nanoparticles (NPs) induce pain resolution, macrophage polarization, and fibrinogen clearance after CCI. **(A)** Treatment plan for RvD1-NPs in CCI-operated male rats: Animals received a perineurial injection of RvD1-NPs or empty NPs. Animals were retested at baseline, 1, 2, 3, and 6 h from days 18 to 24, and sacrificed on day 25. Blue arrows indicate daily injection/test program. T: reflexive testing; syringe: injection. **(B)** Estimated amount of released RvD1 from NPs by an dialysis experiment. The NP solution was diluted in PBS and incubated at 37°C in dialysis tubes. The RvD1 concentration of the dialysates were subsequently determined by LC-MSMS (n = 1). **(C)** Mechanical allodynia was measured after the first RvD1-NP injection on day 18 (n = 6, two-way repeated measures ANOVA with Tukey’s multiple comparisons). **(D)** Mechanical hypersensitivities measured before the injections during the injection period (n = 6, two-way repeated measures ANOVA with Tukey’s multiple comparisons). **(E)** Representative images of fibrinogen immunostaining in sciatic nerve cross sections after RvD1-NP injections. The dashed lines indicate the endoneurial region. Scale bar: 300 µm. **(F)** Quantification of the intensity of fibrinogen immunoreactivity within the endoneurial regions (n = 6, Student’s t-tests with Welch’s correction). **(G-K)** Relative mRNA expression of *Tnfα*, *Arg1*, *Cd206*, *Nlrp3*, and *Plat* after daily RvD1-NP injections (n = 6, Student’s t-tests with Welch’s correction). **(L)** Fluxes of FITC-labelled fibrinogen through monolayers of primary human dermal microvascular cells (HDMECs) was measured *in vitro*. Extracelluar FITC-fibrinogen fluorescent intensity was assessed to evaluate barrier integrity while intracellular FITC-fibrinogen was assessed to evaluate endocytosis (n=7-9, one-way ANOVA with multiple comparisons). NP: empty nanoparticles; RvD1-NP: RvD1-laden nanoparticles. Scale bar: 300µm. All data are shown as mean ± SEM, *: p<0.05.

To assess the effects on the endothelial barrier, we conducted experiments employing primary human dermal microvascular cells (HDMECs), which are of endothelial origin and possess typical features that are present in endoneurial vessels as well. Fluxes of FITC-labelled fibrinogen through a monolayer of HDMEC were assessed to study the barrier after challenge with TNFα and treatment with RvD1. Treatment with TNFα significantly increased the paracellular permeability for FITC-fibrinogen, while transcellular routes remained unaffected (**Fig. 7L**). RvD1 was able to completely restore this TNFα-induced barrier defect **(Fig. 7L)**.

### 7. Fpr2-fibrinogen clearance pathway is inhibited by TAM receptor blocking

Schwann cells are the main drivers of fibrinogen clearance through tPA secretion (Akassoglou et al., 2000; Pellegatta et al., 2022) but do not express Fpr2. Growth arrest-specific 6 (Gas6), the cognate glycoprotein ligand for TAM receptors secreted by macrophages, is necessary for Schwann cell maturation after nerve injury (Stratton et al., 2018). Thus, we hypothesized that Fpr2 activation induces Gas6 release from macrophages, which triggers tPA secretion in Schwann cells and leads to fibrinogen clearance (**Fig. 8A**). Functional evidence supported the TAM pathway: co-injection of the TAM inhibitor RU-301 to block signalling before each BML-111 injection inhibited the analgesic effects of BML-111 on mechanical and thermal hypersensitivity (**Fig. 8B, C**). Under TAM blockade conditions, endoneurial fibrinogen was not cleared although *Plat* levels were not reduced (**Fig. 8D-F**). *Cd206* was not upregulated (**Fig. 8G**). We expected that *Cd206* would not be changed by TAM blocking, since we hypothesized that the *Cd206* and macrophage polarization increase was mainly caused by Fpr2 stimulation. But TAM receptors also mediate macrophage phenotype shift (Vago et al., 2021) and are also necessary for the Fpr2-mediated macrophage phenotype shift. These results indicate that TAMs and their ligands are critically involved in pain resolution and fibrinolysis.

**Figure 8.**
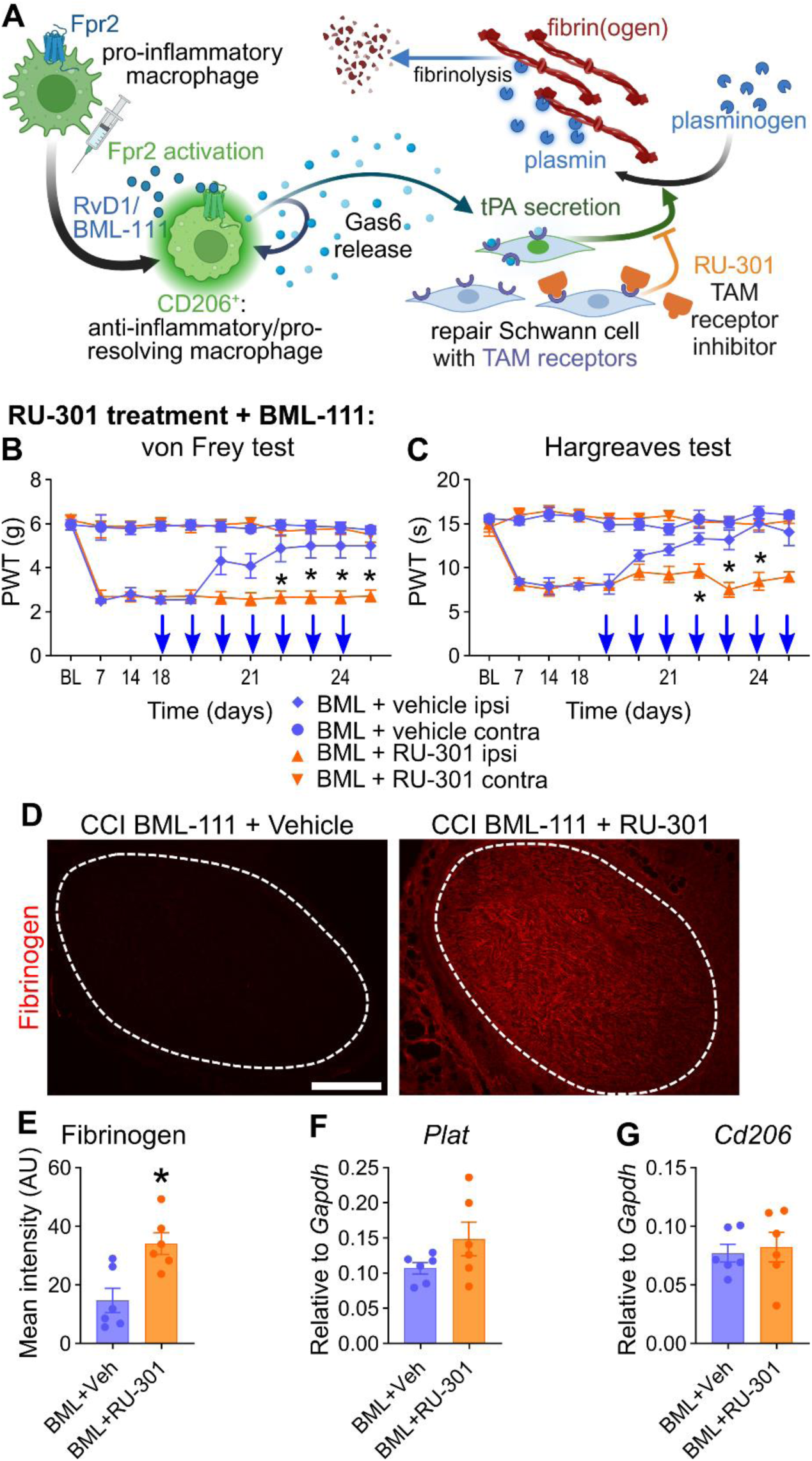
Blocking TAM receptors inhibits Fpr2-mediated pain resolution and fibrinogen clearance. **(A)** Proposed model of pain relief through Fpr2 activation and involvement of the TAM receptor family of receptor tyrosine kinases (Tyro3, Axl, and MerTK) in macrophages and Schwann cells leading to fibrinogen clearance: For details see text. **(B, C)** Male Wistar rats received BML-111 injections combined with prior injections of the TAM receptor inhibitor RU-301 for one week. Paw withdrawal thresholds for **(B)** mechanical and **(C)** thermal hypersensitivities (n= 6, two-way repeated measures ANOVA with Tukey’s multiple comparisons). **(D, E)** Representative images of fibrinogen immunostaining and quantification within the endoneurial regions in sciatic nerve cross sections after BML-111 and/or RU-301 injections. The dashed lines indicate the endoneurial region (n = 6, Student’s t-tests with Welch’s correction). Scale bar: 300 µm. **(F, G)** Relative mRNA expression of *Cd206* and *Plat* after one week BML-111 with and without TAM blocking.

## Discussion

In this study, we characterized the pathophysiology of the injured sciatic nerve at the switch towards pain resolution and identified local treatment options for its acceleration. Beyond the known pathways, we included neural barriers, the coagulation system, and TAM receptors in this process. Pain resolution and functional recovery were independent processes in both male and female rats. Hyperalgesia was intertwined with enhanced diffusion and failed removal of endoneurial fibrinogen, whereas all other neural barriers were intact or persistently open. Using a translational approach, we further showed that – in line with the findings of the rat model of neuropathy – fibrinogen immunoreactivity was enhanced in nerve biopsies from neuropathic patients with pain in comparison with those without pain. Lipidomic screening of SPMs and their receptors identified Fpr2 ligand precursors as switch points. Fpr2 ligands, such as BML-111 and RvD1-NPs, profoundly boosted fibrinogen cleasrance and pain resolution beyond immediate application. TAM receptors mediate this signalling cascade through fibrinolysis. In patients with painful polyneuropathies, endoneurial fibrinogen deposition and anti-inflammatory activity are increased, which might indicate similar mechanisms. These novel findings provide new options for the treatment of unresolved neuropathic pain when fibrinogen deposition is prevalent.

To map the spatiotemporal pattern to specific behavioural recovery patterns, we conducted a battery of reflexive and non-reflexive behavioural tests and, in parallel, permeability assays to correlate barrier recovery with pain resolution 3 to 6 weeks after CCI. The course of **thermal and mechanical allodynia** after CCI in male and female rats was similar to that previously described (Sommer et al., 1993). Gait patterns did not completely recover, which might be attributed to residual muscle weakness or coordination deficits, and possibly higher sensitivity compared to von Frey in CCI rats (Chiang et al., 2014). However, CCI rats still reached similar distances and speeds on the voluntary running wheel, suggesting that their overall activity was unaffected.

In our **permeability assays**, none of the nerve barriers completely recovered with the resolution of pain. The **perineurium** remained leaky until week 9, which was also reflected by the low expression of tight junction barrier proteins. Similar findings have been previously reported (Hirasawa et al., 1994). In our **myelin barrier** analysis, we used the non-degenerating proximal nerve for immersion, but the ligation part for *Cldn19*, which would easily explain the conflicting results. Interestingly, the **capillary barrier** remained permeable to the EBA tracer until 6 weeks; however, at this time point, fibrinogen deposition was already cleared from the endoneurium. We postulate that **fibrinogen clearance from the endoneurium is necessary for complete recovery of the capillary barrier**. Endoneurial fibrinogen drives axonal degeneration and prevents Schwann cell redifferentiation (Akassoglou et al., 2000; Akassoglou et al., 2002; Davalos & Akassoglou, 2012), while successful clearance of fibrinogen and myelin debris allows Schwann cell remyelination (Akassoglou et al., 2002; Pellegatta et al., 2022). In turn, dedifferentiated Schwann cells induce opening of the capillary barrier, as evidenced by immune cell infiltration and EBA tracer extravasation, both of which recover when Schwann cells redifferentiate (Napoli et al., 2012), indicating that barrier closure is dependent on fibrinogen clearance. This could explain the delayed complete recovery of the EBA tracer permeability. The tight junction mRNA expression was in accordance with the results of the permeability assay. However, *Malong et al.* showed in a non-injury model that Schwann cells exclusively induce barrier breakdown by increasing transcytosis rates, while tight junctions remained intact (Malong et al., 2023). The exact mechanism of leakage remains unknown in our analysis. Pain **resolution coincided with the clearance of endoneurial fibrinogen** rather than with inflammation or resealing of capillary vessels for smaller molecules. As **fibrinogen** is a pro-nociceptive and pro-inflammatory molecule (Lim et al., 2014), **its degradation,** together with less diffusion, alleviates hypersensitivity.

In our **lipidomic analysis**, only 11 out of 34 lipids were detected in the sciatic nerve of CCI or sham animals. SPMs themselves were undetectable. This may be because of nutritional or technical reasons. First, the standard chow is probably not rich in omega-3 acids. Indeed, feeding fish oil drastically increases RvD1 levels (Lobo et al., 2016). Second, the methods to measure endogenous SPMs and their detectability in the context of their instability and low concentrations are controversial (Schebb et al., 2022). Exogenously-applied SPMs and their analogs have shown pro-resolving effects in different tissue types and conditions in numerous studies, including ours (Poblete et al., 2022; Tao et al., 2020). Therefore, we decided to concentrate on the precursors and chose **AT-LXA4 for binding to the FRP2** receptor. Locally-applied Fpr2 ligands (BML-111 and RvD1-nanoparticles) alleviated thermal and mechanical allodynia and cleared endoneurial fibrinogen independent of sex. In fact, the encapsulation of SPMs into nanoparticles seems to be a promising technology for the delivery of anti-inflammatory substances (Miranda et al., 2023; M. Quiros et al., 2020).

How does Fpr2 activation induce fibrinolysis? Schwann cells are the primary drivers of **fibrinolysis** after nerve injury via the release of tPA (Akassoglou et al., 2000). However, Fpr2 is mainly found in macrophages and not Schwann cells. Indeed, our data showed elevated *Plat* levels after activation of the Fpr2 receptor. As in previous studies, Fpr2 activation also increased *Cd206 levels*, indicating a macrophage shift towards an anti-inflammatory/pro-resolving phenotype. However, the link between these two macrophages and tPA-producing Schwann cells remains unknown. We hypothesized that macrophages are triggered to release Gas6, which has been shown to induce the maturation of Schwann cells (Stratton et al., 2018), and that TAM receptor signalling is involved in this process. We demonstrated that TAM receptors play a crucial role in preventing pain resolution and fibrinogen clearance.

The effect of Fpr2 activation is most likely multifactorial and reciprocal. In our *in vitro* barrier model, RvD1 protected the monolayer against fibrinogen flux after TNFα challenge. But also TAM signalling and fibrinogen act on endothelial cells (Avanzi et al., 1998; Happonen et al., 2016; Muradashvili et al., 2014). Fibrinogen increases endothelial permeability (Patibandla et al., 2009; Paul et al., 2007; Tyagi et al., 2008), whereas TAM signalling strengthens barrier integrity (Miner et al., 2015). **The role of macrophages as guardians of the capillary barrier** involves uptake of infiltrating molecules. Under homeostatic conditions, the endothelial barrier in the nerve and dorsal root ganglion allows minimal amounts of blood molecules to pass through, which are taken up by macrophages (Lund et al., 2024; Malong et al., 2023). TAMs can shift macrophages to a phenotype with pro-resolving properties (Vago et al., 2021). Fpr2 activation might lead to pain resolution through the macrophage phenotype shift, which further activates TAM signalling, leading to sustained *Cd206*-expressing macrophage population, fibrinolysis, plasmin synthesis, debris removal, and BNB resealing, alleviating neuroinflammation and thereby neuropathic pain.

**The strengths** of this study include the diverse array of experimental approaches used to examine the mechanical and thermal hyperalgesia in male and female rats. Furthermore, we examined fully-establisheded neuropathic pain and developed a suitable intervention option not only for prevention. We provide evidence that neuropathic pain is triggered by fibrinogen deposition both in an animal model of and in humans with neuropathic pain. We provided a promising treatment option (Fpr2 activation) and an optimized delivery system (RvD1-laden nanoparticles) to resolve neuropathic pain. In addition, we investigated the potential downstream therapeutic targets for therapeutic intervention (TAM receptors and fibrinolysis). **The limitations** of this study include missing follow-up investigations of patients. We did not elucidate whether the patients’ pain resolved and lacked healthy controls. Furthermore, the patients were relatively old and did not suffer from nerve injury; therefore, different mechanisms might apply. Human nerve biopsies are rarely performed, and regulations are strictly defined. Therefore, we could not restrict patient inclusion to stringent criteria. We only investigated the CCI model. While similar mechanisms could apply to other models with Wallerian degeneration such as nerve crush or spared nerve injuries, this might not be translatable to other neuropathic pain conditions such metabolic or hereditary neuropathies. Another limitation was the underlying role of fibrinogen in the maintenance of hypersensitivity. Our study lacks direct proof that fibrinogen causes neuropathic pain in the rat CCI model. Conceptually, direct and conclusive evidence for the role of fibrinogen in neuropathic pain could be obtained by examining fibrinogen deficient mice (Drew et al., 2001) or by lipid nanoparticle encapsulated siRNA mediated suppression of fibrinogen production (Juang et al., 2022). In a translational approach, fibrinogen deposition was elevated in nerve biopsies from patients with painful vs. non-painful neuropathy but, again, this evidence is not conclusive.

The mechanistic pathway presented in this study serves as the basis for the development of therapeutic targets for neuropathic pain. The regulation of macrophage phenotypes and coagulation factors by SPMs and TAM ligands might be a promising approach for inflammatory polyneuropathies in which macrophages accumulate. Tackling the local effects of fully-developed neuropathic pain has the potential to resolve chronic neuropathic pain, as it boosts the pain resolution switch. The use of nanoparticles as a delivery system further enables the application of short-lived substances and broadens the molecular toolbox for patient treatment.

## Conclusion

This study linked neuropathic pain after nerve injury with capillary leakage in male and female rats. Both pain and diffused endoneurial fibrinogen subsided after local intervention with Fpr2 ligands delivered by nanoparticles. The functional pathway includes a anti-inflammatory/pro-resolving macrophage shift and TAM receptor signalling. This novel concept in the field of pain resolution may shed light on new targets for the treatment of peripheral neuropathies.

## Materials and Methods

### 1. Animal model

All animal experiments were approved by the Government of Lower Franconia, Germany (55.2.2-2532-2-612 – 06.04.2018 and 55.2.2-2532-2-1547 – 25.04.2022, Regierung von Unterfranken). Wistar rats (Janvier Labs, Le Genest-Saint-Isle, France) were maintained under pathogen-free conditions with a controlled light cycle, temperature, and humidity (14:10 h light/dark cycle, 20–24°C, 45–65% humidity). Rats (200–250 g) were provided standard chow and water *ad libitum* and randomly assigned to the experimental groups. For CCI, rats were deeply anaesthetized with 2-4% isoflurane. The right sciatic nerve was surgically exposed, and four loose silk ligatures were placed around it with a spacing of approximately 1 mm. The incision was closed using sutures. Animals with auto-mutilation were excluded from experiments. Sham-operated rats served as the controls. All experiments were performed in accordance with ethical regulations.

### 2. Perineural injection

Perineurial injections were performed under isoflurane anesthesia. A blunt cannula attached to a nerve stimulator was inserted through the skin after pre-puncture with a 22-Gauge canule. When proximity to the nerve was confirmed by foot twitching, 0.3 ml of 500 nmol BML-111/phosphate buffer saline (PBS; Merck KGaA, Darmstadt, Germany; Cat. No.: SML0215-25MG) or 1 mg RU-301/64% DMSO/PBS (Biozol, Cat. No.: MCE-HY-119039) or vehicle was injected. For the injection of the nanoparticles, the volume was 0.1 ml.

### 3. Behavioural testing reflexive and non-reflexive

The von Frey test was performed to assess mechanical allodynia using the Dixon’s up-and- dosswn method. Von Frey filaments (Aesthesio® set, Ugo Basile SRL, Gemonio, Italy) were applied to the plantar surface of each hind paw for 1-3 s, so that the filaments were bent to a 45°-angle. Single paws were tested with a recovery time of at least 30 s between applications. For analysis, the means of the filament forces with positive reactions were averaged per paw. Thermal hypersensitivity was assessed by the Hargreaves test. The light source of a Plantar Analgesia Meter (Model 400 Heated Base from IITC Life Science, Woodland Hills, California, USA) was applied to the plantar surface of each hind paw with a cut-off latency of 20 s. Withdrawal latencies were recorded twice (at least 30 s intervals) and averaged for analysis. Motor activity was assessed by voluntary wheel running (Model BIO-ACTIVW; Bioseb, Vitrolles, France). Once per week, the rats were individually placed in cages with voluntary running wheels for 24 h. The activity parameters, including distance, speed, and acceleration, were recorded using the BIO-ACTIVW-SOFT (Bioseb) software.

The CatWalk XT system (Noldus, Wageningen, Netherlands) was used for gait analysis. The rats were allowed to move freely on the CatWalk until three valid runs were recorded with a camera. Runs with a run duration of 0.5 - 5 s and a maximal run speed variation of 60% were classified as valid. The ratios of footprint area and standing time between the right and left hind paws were calculated. Data from three valid runs were averaged for the analysis.

### 4. Reverse transcription quantitative PCR

To isolate RNA from snap-frozen sciatic nerve tissue (site of ligation in CCI), we used the RNeasy Micro Kit (#74004, Qiagen, Venlo, The Netherlands) according to the manufacturer’s instructions. RNA concentration was determined using a NanoDrop ND 2000 spectrophotometer (Thermo Fisher Scientific). 1000 ng (for the 1–6-week characterization) and 500 ng of total RNA was reverse-transcribed to cDNA using the High-Capacity cDNA Reverse Transcription Kit, according to the manufacturer’s instructions (Thermo Fisher Scientific). For quantitative polymerase chain reaction (qPCR), the following TaqMan gene expression assays with FAM-labeled probes were used (Applied Biosystems, Thermo Fisher Scientific): *Gapdh* (Rn01462662_g1), *Tnfα* (Rn99999017_m1), *Cd68* (Rn01495634_g1), *Plat* (Rn01482578_m1), *Tjp1* (Rn02116071_s1), *Cldn1* (Rn00581740_m1), *Cldn5* (Rn01753146_s1), *Cldn12* (Rn04219013_m1), *Cldn19* (Rn01416537_m1), *Mpz* (Rn00566746_m1), *Gpr18* (Rn01493247_m1), *Gpr37* (Rn00589441_m1), *Gpr37l1* (Rn00595762_m1), *CmklR1/ChemR23* (Rn00573616_s1)*, Fpr2* (Rn03037051_gH), *Lgr6* (Rn01490727_m1) *Arg1* (Rn00691090_m1), *Mrc-1/Cd206* (Rn01487342_m1), *Nlrp3* (Rn04244620_m1). qPCRs were run on a StepOnePlus real-time PCR system (Applied Biosystems) using the following program: 40 cycles at 95°C for 1 s and 60°C for 20 s. The samples were run in triplicate and averaged for analysis. mRNA abundance relative to the reference gene *Gapdh* is presented as 2^-ΔCt^.

### 5. Permeability assessment: perineurial barrier, myelin barrier, endoneurial vessels

To assess perineurial permeability, ligation endings of the sciatic nerve were sealed with Vaseline and immersed in 1 ml of 5% EBA (5% bovine serum albumin, 1% Evans Blue) and 3% sodium fluorescein (NaFlu) in PBS for 15 min, followed by 5% EBA for 45 min at room temperature (RT). After washing with PBS, the nerves were fixed in 4% paraformaldehyde (PFA) for 1-3 h, cryoprotected in 30% sucrose at 4°C overnight, and frozen in Tissue-Tek® O.C.T.™ Compound (Sakura, Alphen aan den Rijn, the Netherlands). 10-µm sections were cut using a cryostat (Leica CM3050S; Leica Biosystems, Wetzlar, Germany). Immunofluorescence images were acquired using a microscope (BZ-9000, Keyence Deutschland GmbH, Neu-Isenburg, Germany) and the intensity was measured.

For myelin barrier assessment, nerves proximal from the ligation were desheathed and sealed with Vaseline at their endings (Chen et al., 2021). After incubation in artificial cerebrospinal fluid (10mM HEPES, 110 mM NaCl, 17.8 mM NaHCO_3_, 4 mM MgSO_4_, 3.9 mM KCl, 3 mM KH_2_PO_4_, 1.2 mM CaCl_2_, 10 mM Dextrose) containing 0.5% 70 kDa fluorescein isothiocyanate-dextran (FITC-Dextran; Sigma-Aldrich, St. Louis, MO, USA) for 1 h at 37°C, washed, fixed in 4% PFA for 5 min at RT, and teased to single nerve fibers on microscopy slides. Until mounting with Aqua-Poly/Mount medium, the slides were stored at -20°C. Fluorescence images were then acquired for analysis.

To assess the permeability of the endoneurial blood vessels, rats received an intravenous injection of 1 ml 5% EBA per 100 g body weight (Omura et al., 2004). After 30 min, the proximal and distal parts of the sciatic nerve were snap-frozen in Tissue-Tek® O.C.T.™ Compound. Immunofluorescence images of 10-µm sections were acquired, and the endoneurial intensity was measured.

### 6. *In vitro* assessment of barrier recovery by resolvin D1 and permeability for fibrinogen

HDMECs (PromoCell, Heidelberg, Germany) were employed as endothelial/endoneurial model and seeded on Millipore filter inserts (Merck-Millipore, PCF-filters, pore-size 0.45 µm, area 0.6 cm²) and grown until confluent and stable in transendothelial resistance (TER). Cells were then challenged with 500 U/ml TNFα (Peprotech) for 24 h in parallel to untreated controls. Then TNFα concentration was reduced to avoid induction of apoptosis to 100 U/ml, and some filters were additionally incubated with 500 nM RvD1 (Cayman Chemicals/Biomol GmbH, Hamburg, Germnay) for further 24 h. Permeability for fibrinogen was determined, using FITC-labelled fibrinogen (LOXO GmbH, Dossenheim, Germany), which was added in a concentration of 0.58 µM apically to the filters. After 4, 5 or 6 h, concentrations of FITC-fibrinogen in the basal compartments were measured fluorometrically (Tecan Infinite M200, Tecan, Switzerland). To analyse potential changes in transcellular passage, e.g. by endocytosis, cells were harvested from the filters after washing with PBS and were dissolved using total lysate buffer (10 mM Tric-Cl pH 7,5, 150 mM NaCl, 0,5% Triton X-100, 0,1% SDS, protease inhibitors). Intracellular concentration of FITC-fibrinogen were detected at the Tecan photometer. Tracer fluxes and apparent permeabilities were calculated from the concentrations of FITC-fibrinogen.

### 7. Liquid Chromatography tandem mass spectrometry

The SPMs were determined as described in a previous study by *Toewe et al.* (Toewe et al., 2018) after some modifications of the extraction method. After dissection, sciatic nerve tissue was snap-frozen in liquid nitrogen. Tissue samples were homogenized in water:ethanol (75:25, v/v) to a tissue homogenate of 0.1 mg/μl using a swing mill (Retsch, Haan, Germany) with five zirconium oxide grinding balls (25 Hz for 2.5 min). 300 μl of the tissue homogenates (in total 30 mg tissue) were spiked with internal standard solution and extracted using solid-phase extraction with Express ABN cartridges and Extrahera (both from Biotage). Afterwards, the sample extracts were measured using an LC-MSMS system comprising an Agilent 1200 LC system and a 5500 QTRAP mass spectrometer (Sciex). Chiral chromatographic separation was achieved using a Lux Amylose-1 column (250 x 4.6 mm, 3 µm, Phenomenex) with water:FA (99.9:0.1, v/v) as solvent A and ACN:MeOH:FA (95:4.9:0.1, v/v/v) as solvent B in gradient elution mode. Data acquisition and evaluation were performed using Analyst 1.6.2 and MultiQuant 3.0 (both from Sciex).

### 8. Schwann cell experiments

Primary Schwann cell cultures from adult rat sciatic nerves were established as previously described (Andersen & Monje, 2018). 100,000 Schwann cells were seeded into 12-well plates in Dulbecco′s Modified Eagle′s medium (DMEM, high glucose; Sigma, FG0445-500ML) containing 10% fetal calf serum, 1% penicillin/streptomycin, and 25 µg/ml gentamycin, and incubated with 250 µM CPT-cAMP (ab120424; Abcam) for 1-5 days to induce differentiation following standard protocols (Monje, 2018). After incubation, the cells were trypsinized and homogenized using QIAshredder (QIAgen, #79654) for RNA isolation using the RNeasy Mini Kit, according to the manufacturer’s instructions (QIAgen, #74104).

### 9. Production of nanoparticles

Nanoparticles were produced as previously described (Miguel Quiros et al., 2020). Briefly, PLGA-PEG-biotin (Nanosoft Polymers; PLGA molecular weight = 10 kDa, PEG molecular weight = 2 kDa) was dissolved in dimethylformamide (100 mg/ml), and 3.76 μg of RvD1 (Cayman Chemical, Ann Arbor, USA; dissolved in ethanol at 0.1 mg/ml) was added to the polymer solution. Next, 500 µL of this polymer-resolvin mixture was added dropwise to 10 mL of PBS. Unloaded control nanoparticles were prepared by dropwise addition of a polymer solution (without resolvin) to PBS. The nanoparticles were stirred for 4-5 h, concentrated by centrifugation using Amicon Ultra-15 centrifugal filter units, and filtered through sterile 0.45-μm syringe filters.

### 10. Dialysis of nanoparticles to measure resolving release

Slide-A-Lyzer^TM^ mini dialysis unit 20K MWCO (Thermo Fisher Scientific, Darmstadt, Germany; ref: 69590) was loaded with 100 µL of RvD1-laden nanoparticles. The devices were placed in 1.1 mL of PBS. The prepared tubes were stirred in a thermomixer at 300 rpm and 37°C for 1, 3, 6 h, 1, 2, and 3 days. The dialysates were stored at -80°C until shipment for analysis by liquid chromatography-tandem mass spectrometry.

### 11. Immunofluorescence of rat nerve tissue

The nerves were snap-frozen in Tissue-Tek® O.C.T.™ Compound. The 10-µm sections were fixed in a -20°C acetone bath. Blocking and permeabilization were performed using PBS containing 0.3% Triton X-100, 0.1% Tween20 and 10% donkey serum for 1 h at RT. The sections were incubated with goat anti-fibrinogen (1:100; antibodies-online GmbH, Aachen, Germany; Cat # ABIN458743) primary antibodies overnight at 4 °C. Suitable secondary antibodies were applied for 1 h at RT: donkey anti-goat IgG Alexa Fluor 594 (1:600; cat. A32758; Thermo Fisher Scientific). Before mounting with Aqua-Poly/Mount medium (Vector Labs, Burlingame, CA, USA), nuclei were stained with DAPI. In this study, we used a polyclonal fibrinogen antibody that binds to various fibrin(ogen) (degradation) products. In the following section, we will use the term fibrinogen for fibrin.

### 12. Human nerve tissue

The human sural nerves were obtained for diagnostic purposes. Nerve material that was left over after diagnostic work-up was used for the study with approval from the Ethical committee of the University of Würzburg (238/17). For patient details, please refer to Supplementary Tables ST1. Sixteen biopsies from patients with neuropathies were obtained for immunofluorescence, freshly frozen in OCT, and stored at -80°C. 10µm-sections were cut. For staining, sections were thawed, dried, and fixed in 4% PFA for 10 min at RT. After washing, sections were blocked with 10% donkey serum, 0.1% TritonX-100 and 0.3% Tween20 in PBS for 1 h at RT. Rabbit anti-tPA (1:100; 10147-1-AP; Proteintech) and sheep anti-fibrinogen (1:200; ab118533; Abcam) antibodies were incubated overnight at 4°C. Suitable secondary antibodies (1:600; anti-rabbit Alexa Fluor 488, Invitrogen, #A32731; anti-sheep Alexa Fluor 647, Invitrogen, #A21448) were incubated for 1 h at RT. Before mounting with Aqua-Poly/Mount medium (Vector Labs), the nuclei were stained with Hoechst33342.

### 13. Image analysis

Nerves from the permeability assays and fibrinogen staining were imaged using a fluorescence microscope (BZ-9000; Keyence Deutschland GmbH, Neu-Isenburg, Germany). Images were further analysed using ImageJ software (Schneider et al., 2012). The endoneurial region of each image was manually selected, and the mean intensity was measured. For each sample, three images were analysed and averaged.

For the teased fibres, brightfield images were acquired to place a 5-µm-diameter circle as the region of interest (ROIs) in the internodal region of the fibres and three ROIs for background subtraction. These ROIs were transferred to their corresponding fluorescence images and the mean intensities of the ROIs were measured. The intensity of the background ROIs was averaged and subtracted from each internodal ROI.

Image acquisition for human nerve tissue was performed using an Axio Imager 2 fluorescence microscope coupled with an Axiocam 506 mono camera (Zeiss, Oberkochen, Germany). For each sample, 2-3 images were analysed and averaged.

### 14. Statistical analysis

Statistical analyses were performed using GraphPad Prism version 9.3.0 Windows (GraphPad Software, San Diego, CA, USA). For animal behavior tests, two-way repeated measures ANOVA with Tukey’s multiple comparison test was performed. The tests performed included Mann-Whitney-U-test as a non-parametric test or Student’s t-test (incl. Welch’s correction for unequal variance), two-way ANOVA + post-hoc test for multiple groups (see figure legends). Statistical significance was set at p < 0.05. All data are expressed as the mean ± SEM.

## Funding and statement of interests

This study was supported by the German Research Foundation Clinical Research Group ResolvePAIN KFO5001 – 42650386, Ri 817/15-1 – 433191715, and SFB 1039/Z01 as well as Evangelisches Studienwerk Villigst and DAAD.

The authors declare that they have no conflict of interest.

## Author contributions

BH and HLR conceptualized this study. HLR and AB secured the research funding. CS and KD provided human nerve samples. AN and PVM provided valuable guidance on experimental setups and protocols. PPK and AJG produced and provided the nanoparticles. DT, RG, and MS performed the LCMSMS. BH, ABK, SK, CG, IK, and SMK performed experiments. BH, ABK, SK, CG, and SMK analysed data. BH drafted the original manuscript. HLR critically reviewed the first draft of the manuscript for intellectual content. All authors read, commented on, and approved the final manuscript.

## Data availability

The data are available from the corresponding author upon reasonable request.

## Acknowledgments

The authors thank In-Fah M. Lee (Clinical Physiology/Nutritional Medicine, Charité Berlin) for excellent technical assistance.

Schematic illustrations were created with BioRender.com.

The authors have no conflicting financial interests.

---

**Supplementary Figure 1:**
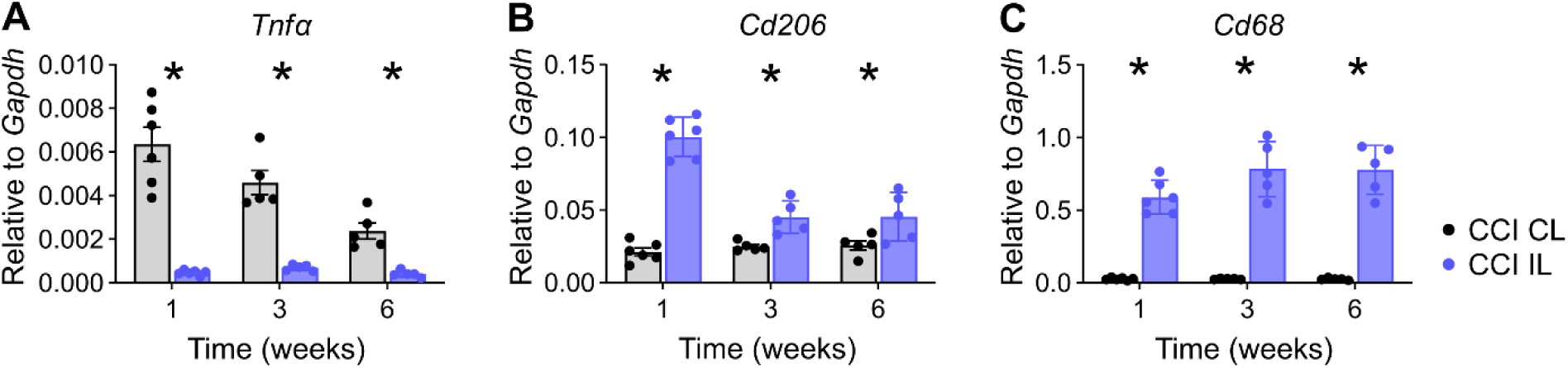
Neuropathic pain resolves at 6 weeks after nerve injury in female rats. Female Wistar rats underwent CCI or sham surgery and were examined weekly. **(A)** Thermal and (**B)** mechanical hypersensitivity were assessed (n = 8). Data in graphs are shown as mean ± SEM, * p<0.05. Two-way repeated measures ANOVA with Tukey’s multiple comparison.

**Fig. S2.**
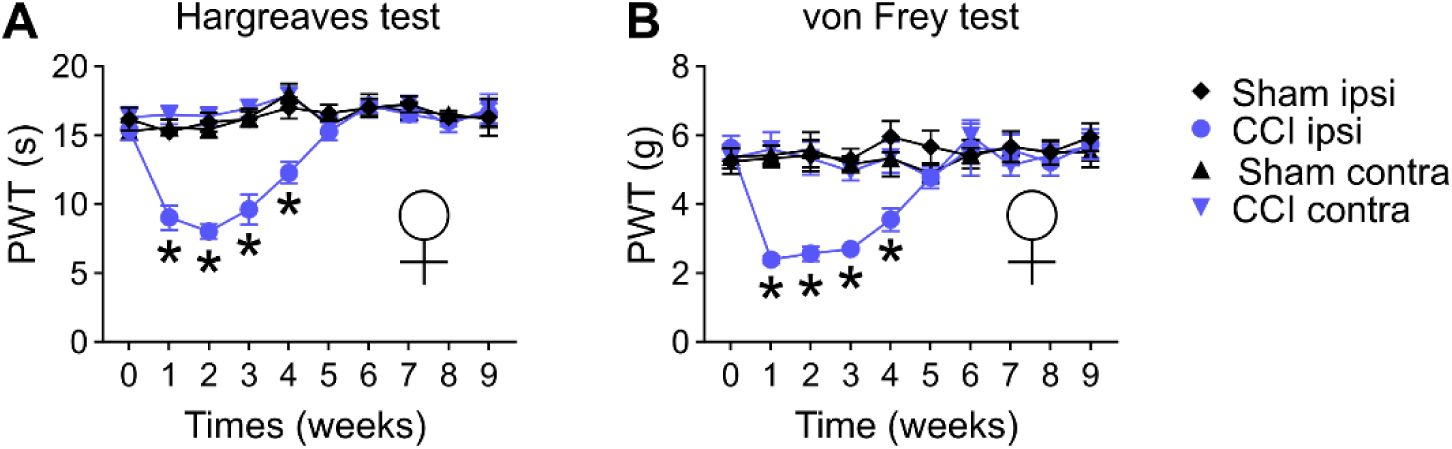
mRNA expression of *Cldn12* and *Mpz* is reduced after nerve injury. The ligation part of ipsilateral (IL) and the contralateral (CL) sciatic nerve was assessed (n = 5-6). All data are shown as mean ± SEM, * p<0.05 compared to control at the indicated time points, two-way ANOVA with Šidák’s multiple comparisons.

**Fig S3:**
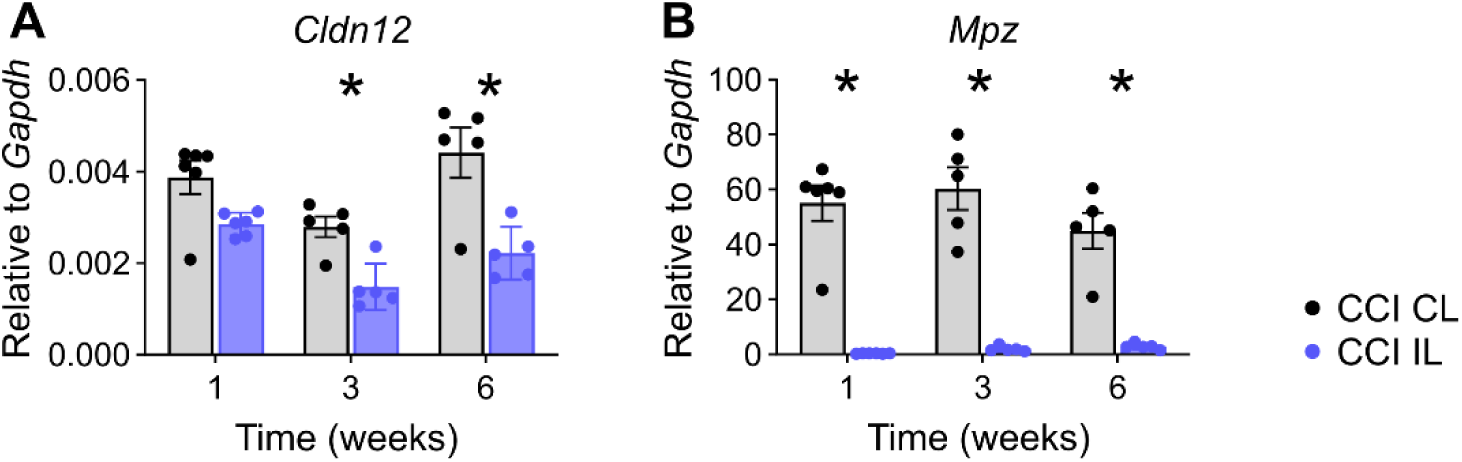
Rapid upregulation of *Gpr37* in Schwann cell cultures after stimulation of differentiation. Primary Schwann cells from adults rat were *in vitro* stimulated with cAMP to induce redifferentiation. **(A)** *Gpr37*, **(B)** *Gpr37l1* and **(C)** *Mpz* mRNA were assessed at designated timepoints. All data are shown as mean ± SEM, n = 5-6, two-way ANOVA with Šidák’s multiple comparisons. * p<0.05.

**Fig S4:**
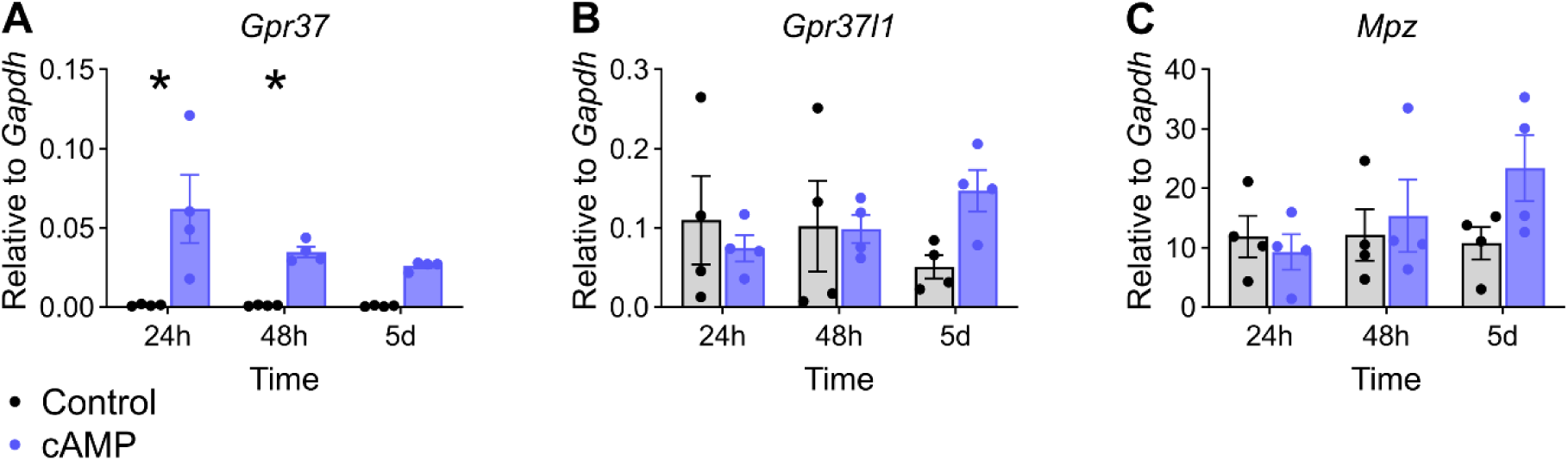
Nerve injury promotes long-term upregulation of *Tnfα* and macrophages markers. The ligation part of ipsilateral (IL) and the contralateral (CL) sciatic nerve was assessed (n = 5-6). All data are shown as mean ± SEM, * p<0.05 compared to control at the indicated time points, two-way ANOVA with Šidák’s multiple comparisons.

**Supplementary Figure 5.**
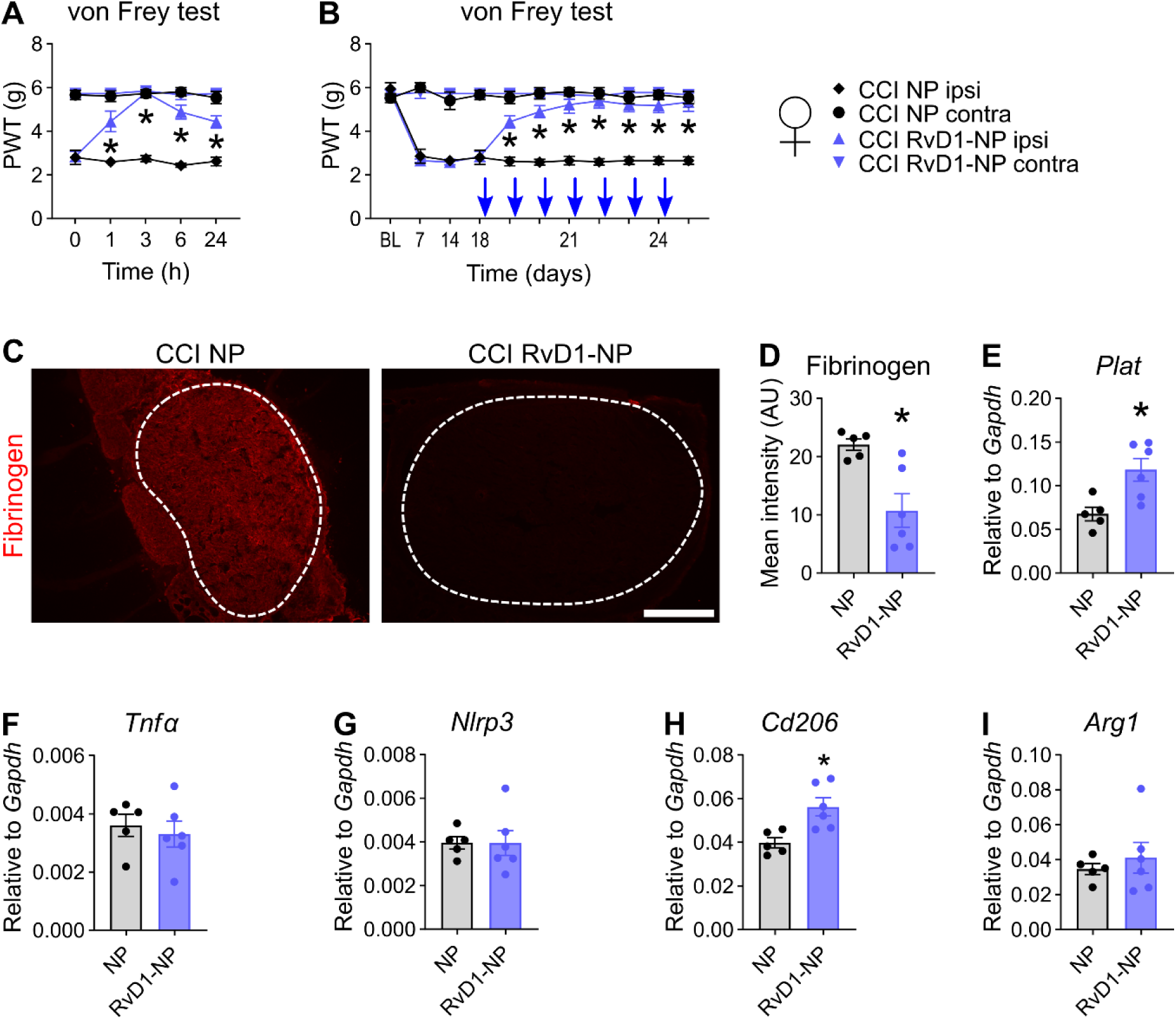
Similar efficacy and effects for RvD1-laden nanoparticles in female rats. On day 18 after CCI, animals received a perineurial injection of RvD1- nanoparticles (NPs) or empty NPs. Animals were retested at 1, 2, 3, and 6 h. This daily injection and test program was performed until day 24. Animals were sacrificed for tissue analysis on day 25. **(A)** Mechanical allodynia was measured after the first RvD1-laden nanoparticle injection on day 18 (n = 6, two-way repeated measures ANOVA with Tukey’s multiple comparisons). **(B)** Mechanical hypersensitivities measured before the injections during the injection period (n = 6, two-way repeated measures ANOVA with Tukey’s multiple comparisons). **(C)** Representative images of fibrinogen immunostaining in sciatic nerve cross sections after RvD1-nanoparticle injections. The dashed lines indicate the endoneurial region. Scale bar: 300µm. **(D)** Quantification of the intensity of fibrinogen immunoreactivity within the endoneurial regions (n = 6, Student’s t-tests with Welch’s correction). **(E-I)** Relative mRNA expression of *Plat*, *Tnfα*, *Nlro31*, *Cd206*, and *Arg1* after daily RvD1-nanoparticle injections (n = 6, Student’s t-tests with Welch’s correction). NP: empty nanoparticles; RvD1-NP: RvD1-laden nanoparticles. All data are shown as mean ± SEM, *: p<0.05.

## Notes

### Competing Interest Statement

The authors have declared no competing interest.

